# Comprehensive Annotations of Human Herpesvirus 6A and 6B Genomes Reveal Novel and Conserved Genomic Features

**DOI:** 10.1101/730028

**Authors:** Yaara Finkel, Dominik Schmiedel, Julie Tai-Schmiedel, Aharon Nachshon, Michal Schwartz, Ofer Mandelboim, Noam Stern-Ginossar

## Abstract

Human herpesvirus 6 (HHV-6) A and B are highly ubiquitous betaherpesviruses, infecting the majority of the human population. Like other herpesviruses, they encompass large genomes and our understanding of their protein coding potential is far from complete. Here we employ ribosome profiling and systematic transcript analysis to experimentally define the HHV-6 translation products and to follow their temporal expression. We identify hundreds of new open reading frames (ORFs), including many upstream ORFs (uORFs) and internal ORFs (iORFs), generating a complete unbiased atlas of HHV-6 proteome. Furthermore, by integrating systematic data from the prototypic betaherpesvirus, human cytomegalovirus, we uncover numerous uORFs and iORFs that are conserved across betaherpesviruses and we show that uORFs are specifically enriched in late viral genes. Using our transcriptome measurements, we identified three highly abundant HHV-6 encoded long non-coding RNAs (lncRNAs), one of which generates a non-polyadenylated stable intron that appears to be a conserved feature of betaherpesviruses. Overall, our work reveals the complexity of HHV-6 genomes and highlights novel features that are conserved between betaherpesviruses, providing a rich resource for future functional studies.

## Introduction

Human herpesvirus 6 (HHV-6) is a highly ubiquitous betaherpesvirus, infecting more than 90% of the human population (Zerr et al., 2005). Based on distinct molecular, epidemiological and biological properties, these viruses were declared as two separate viral species; HHV-6A and HHV-6B (Ablashi et al., 2014). While HHV-6A was not found to be etiologically associated with disease (Braun et al., 1997), HHV-6B was linked to Roseola, leading to febrile seizures in more than 10% of acute infections (De Bolle et al., 2005; Hall et al., 1994; Yamanishi et al., 1988). Both HHV-6A and HHV-6B, like all herpesviruses, establish a lifelong latent infection in their hosts (Kondo et al., 1991; Luppi et al., 1999). HHV-6 latency is established in multiple cell types, where the viral genome is integrated into subtelomeric chromosome regions (Arbuckle et al., 2013; Braun et al., 1997). Remarkably, in approximately 1% of the population worldwide HHV-6 is integrated in every cell in the body, and inherited, due to integration of the viral genome in germline cells (Clark, 2016; Pellett et al., 2012). HHV-6 reactivation is a common cause of encephalitis, and has been associated with several diseases including multiple sclerosis, hepatitis, pneumonitis and graft-versus-host disease (Braun et al., 1997; Caselli and Luca, 2007; De Bolle et al., 2005).

The genomes of HHV-6A and HHV-6B, similar to those of other herpesviruses, consist of large linear double stranded DNA molecules, 143kb in length, containing a unique segment flanked by direct repeats (Lindquester and Pellett, 1991; Martin et al., 1991). The annotation of HHV-6 coding capacity has traditionally relied on open reading frame (ORF)-based analyses using canonical translational start and stop sequences and arbitrary size restriction to demarcate putative protein-coding genes, resulting in a list of around 100 ORFs for each virus (Dominguez et al., 1999; Gompels et al., 1995; Gravel et al., 2013). In recent years, genome wide analysis of herpesviruses using short RNA sequencing (RNA-seq) reads, and recently also direct and long-read RNA-seq revealed very complex transcriptomes (Balázs et al., 2017; Depledge et al., 2019; Gatherer et al., 2011; Kara et al., 2019; O’Grady et al., 2019, 2016; Tombácz et al., 2017), and combined with genome-wide mapping of translation, revealed hundreds of new viral ORFs (Arias et al., 2014; Bencun et al., 2018; Stern-Ginossar et al., 2012; Whisnant et al., 2019). Specifically for HHV-6, recent work using proteomics, transcriptomics and comparative genomics on HHV-6B enabled re-annotation of several viral gene products (Greninger et al., 2018). Taken together, this unforeseen complexity of herpesviruses suggests the current annotations of HHV-6 genomes are likely incomplete.

Here, we apply ribosome profiling (Ribo-seq) and RNA-seq to investigate the genomes of the closely related HHV-6A and HHV-6B. These powerful tools allowed us to accurately determine the translation initiation sites of previously annotated genes, and to identify hundreds of new open reading frames including many upstream ORFs (uORFs) and internal ORFs (iORFs), generating a complete atlas of HHV-6 translation products. Using our RNA-seq data, we were able to map novel splice junctions and to identify novel highly abundant viral long non-coding RNAs. The systematic annotations of two betaherpesviruses together with our previous annotation of the prototypic betaherpesvirus human cytomegalovirus (HCMV)(Stern-Ginossar et al., 2012) created for the first time an opportunity to look at functional conservation of some of these features. We found high levels of conservation between HHV-6A and HHV-6B, and in several cases the newly identified features were also conserved in HCMV. Our results shed light on the complexity of herpesviruses, point to conserved features and can serve as a valuable resource for future studies of these important viruses.

## Results

### Profiling the transcriptome and translatome of HHV-6A and HHV-6B

To capture the full complexity of HHV-6A and HHV-6B genomes, we applied next generation sequencing methods that map genome-wide RNA expression and translation to HSB-2 and Molt-3 cells infected for 72 hours with HHV-6A strain GS and HHV-6B strain Z29, respectively (Figure 1A). For each virus we mapped genome-wide translation events by preparing three different ribosome-profiling libraries (Ribo-seq). Two Ribo-seq libraries facilitate mapping of translation initiation sites, by treating cells with lactimidomycin (LTM) or harringtonine (Harr), drugs that inhibit translation initiation in distinct mechanisms and lead to accumulation of ribosomes at translation initiation sites [Figure 1A and (Ingolia et al., 2011; Lee et al., 2012)]. The third Ribo-seq library was prepared from cells treated with the translation elongation inhibitor cycloheximide (CHX), and gives a snap-shot of actively translating ribosomes across the body of the translated ORF (Figure 1A). In parallel, we used a tailored RNA-sequencing (RNA-seq) protocol which on top of quantification of RNA levels allows identification of transcription start sites (TSSs) due to a strong overrepresentation of fragments that start at the 5’ end of transcripts (Stern-Ginossar et al., 2012), as well as detection of polyadenylation sites (Figure 1A). The combination of these methods provides accurate mapping of transcription and translation events, as seen in the example of U54 (Figure 1A). The different Ribo-seq libraries generate distinct profiles across the coding region, displaying a strong peak at the translation initiation site, which as expected is more distinct in the Harr and LTM libraries, while the CHX library provides the distribution of ribosomes across the entire coding region up to the stop codon. These profiles were consistent across coding regions in human genes (Figure 1B) and, as expected, the RNA-seq profiles were uniformly distributed across the coding region (Figure 1C).

**Figure 1.**
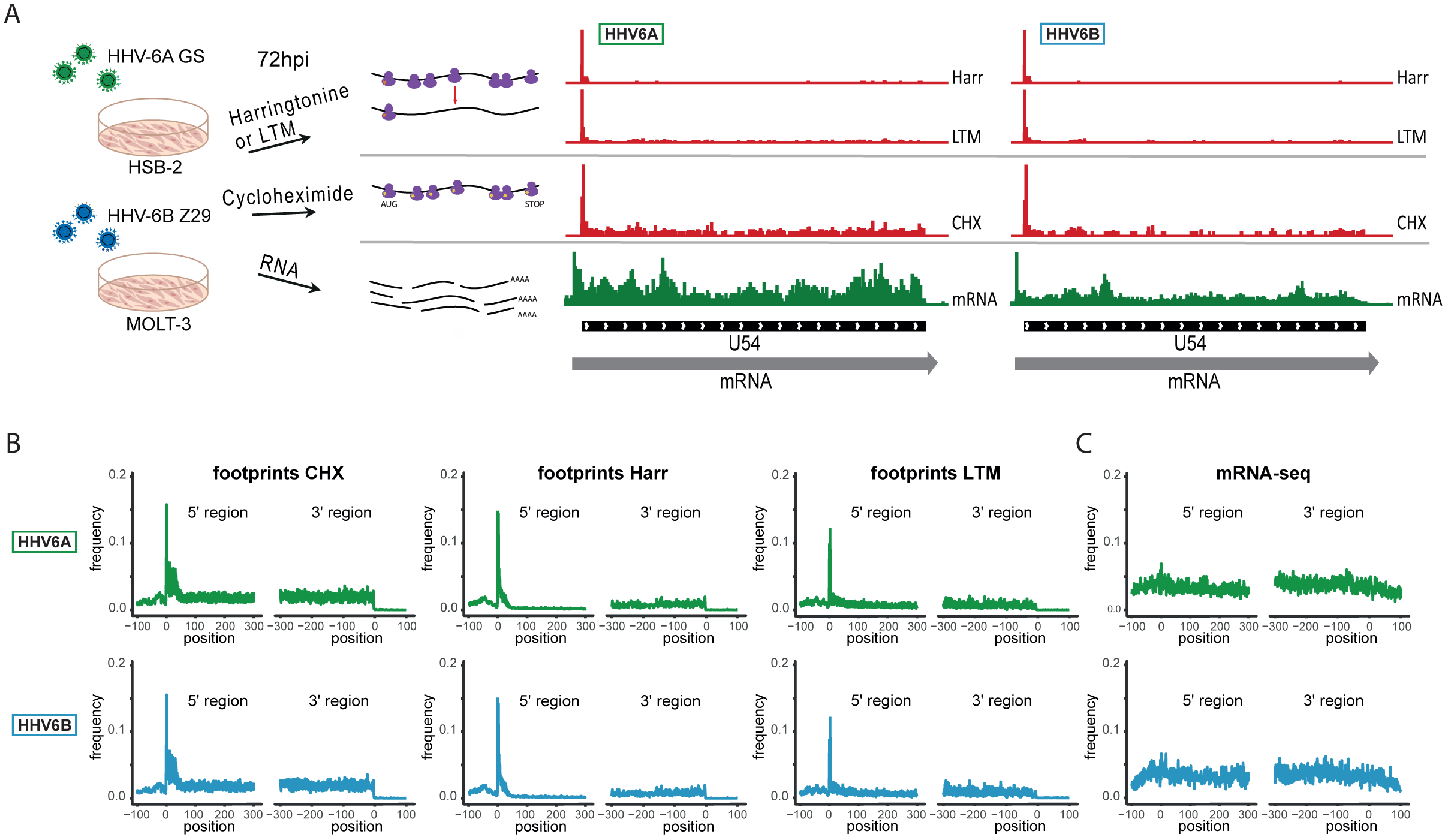
Overview of the experimental approach. (A) Viral gene expression was analyzed by performing ribosome profiling (red) and initiation enriched RNA-seq (green). HSB-2 cells were infected with HHV-6A strain GS, and MOLT3 cells were infected with HHV-6B strain Z29. Infected cells were harvested at 72 hours post infection (hpi) for RNA-seq, and for ribosome profiling using cycloheximide (CHX) treatment to map overall translation or lactimidomycin (LTM) and Harringtonine (Harr) treatments for mapping translation initiation. (B) Metagene analysis of the 5’ and the 3’ regions of human protein coding mRNAs showing the expression profile as measured by the different (B) Ribo-seq and (C) RNA-seq methods in HHV-6A (green) and HHV-6B (blue). The X axis shows the nucleotide position relative to the start or the stop codons.

### Ribo-seq libraries uncover the translation landscape of HHV-6A and HHV-6B

We used the Ribo-seq data to determine translation of viral ORFs. Comparing to previously annotated ORFs, we found many misannotations and novel un-annotated ORFs. Importantly, many of these new ORFs are conserved between HHV-6A and HHV-6B, validating our approach, and emphasizing the high similarity between these two viruses. One example of misannotation is the U30 gene, an essential viral gene coding for an inner tegument protein (Nicholas and Martin, 1994). We found translation of this gene to initiate at an ATG 411bp downstream of the previously annotated start, in both HHV-6A and HHV-6B, resulting in a 946 amino acid (aa) long protein (Figure 2A). Importantly, the new annotations include the C-terminal domain which was shown to interact with the large tegument protein in the HSV-1 homolog (Richards et al., 2017).

**Figure 2.**
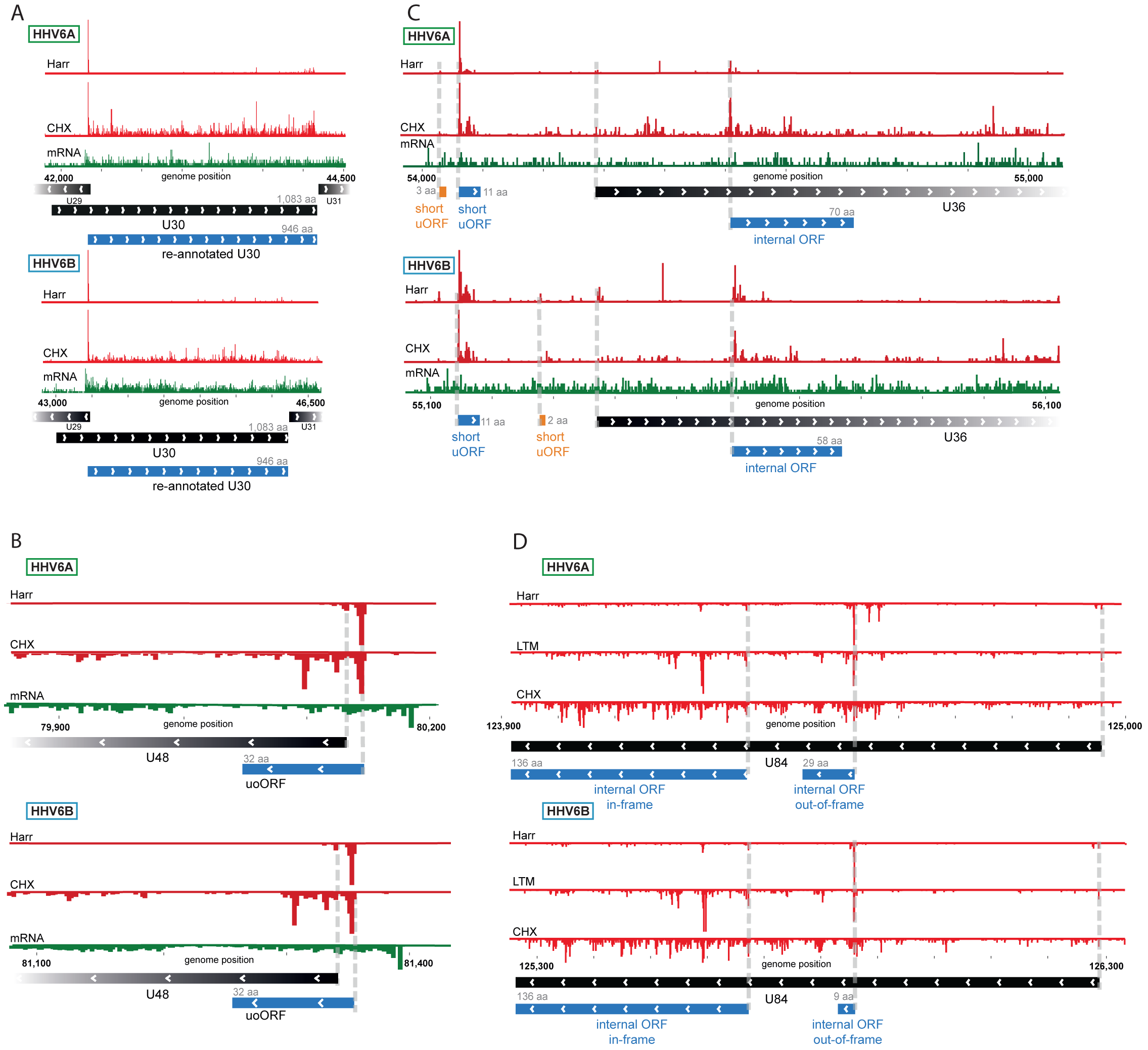
Ribo-Seq measurements reveal the architecture of viral coding regions. Examples of expression profiles of viral genes that contain novel ORFs conserved in HHV-6A and HHV-6B. Ribo-seq reads are presented in red and RNA-seq reads are presented in green. Canonical annotated ORFs are labeled by black rectangles, novel ORFs initiating at an AUG codon are labeled in blue, and novel ORFs initiating at a near-cognate start codon are labeled in orange. ORF sizes are written in grey. LTM profiles are not presented when they fully correspond to the Harr profile. (A) U30 translation initiates exclusively at an AUG downstream of the annotated start codon. (B) A 32 amino acid (aa) upstream overlapping ORF (uoORF) is coded by the U48 transcript, initiates upstream of the U48 canonical ORF and partially overlaps it. (C) U36 locus contains two uORFs, as well as an out-of-frame iORF. (D) U84 locus contains an in-frame iORF which is a truncated version of U84, and a novel out-of-frame iORF.

We identified novel ORFs that are present in both viruses. For example, a short 32aa ORF was found to initiate upstream of the envelope protein gene U48 (Figure 2B). This ORF partially overlaps the U48 gene, making it an upstream overlapping ORF (uoORF). Since uoORFs are known to have repressive regulatory effects conserved across vertebrates (Johnstone et al., 2016), this novel ORF likely negatively regulates the translation of U48. We did not observe translation of another downstream ORF that could be positively regulated by this uoORF. The packaging gene U36 is an example of a gene for which we found translation of two very short (<20aa) uORFs from its 5’UTR (Figure 2C). In addition, we identified translation of an internal ORF (iORF), initiating out-of-frame, inside the coding region of U36 (Figure 2C), leading to translation of a novel ORF. In the U84 gene we observed two iORFs, one of them out-of-frame possibly regulating the downstream ORF, and another in-frame, starting at an ATG downstream of the U84 start-codon and ending in the same stop-codon, resulting in a truncated version of U84 (Figure 2D).

### RNA-seq analysis reveals pervasive splicing that is conserved between HHV-6A and HHV-6B

To systematically map the splice junctions of HHV-6A and HHV-6B we used two independent splice-aware alignment tools, TopHat (Trapnell et al., 2009) and STAR (Dobin et al., 2013). We found an intricate set of splice junctions including dozens of novel splice junctions, which were overall positionally conserved between HHV-6A and HHV-6B (Figure 3A and table S1 and S2). We were able to detect 24 out of 26 annotated HHV-6A splice junctions and all 24 annotated HHV-6B splice junctions. Furthermore, we identified 37 novel splice junctions in HHV-6A and 44 in HHV-6B (Figure 3B and table S1 and S2). Some of the novel splice junctions identified in HHV-6A were recently reported in HHV-6B and are confirmed here for both viruses (U19, U83, and all splice forms of U79) (Greninger et al., 2018). Interestingly, many of the novel splice junctions seem to belong to one long transcript composed of several short exons separated by long introns, spanning the U42-U57 locus (Figure 3A). In a few cases, novel splice junctions result in reannotation of ORFs. For example, a splice junction between the HHV-6B U7 and U8 indicates that they are fused to one translation product, similar to the HHV-6A U7 (Dominguez et al., 1999; Gompels et al., 1995; Gravel et al., 2013) (Figure S1A). Another splice junction in the HHV-6A U13 gene indicates that the U12 and U13 proteins share their N-terminal domain (Figure S1B). The same junction was also detected at lower levels in HHV-6B. The high relative abundance of reads that capture splice junctions suggests there is an extensive use of alternative splicing in these viruses.

**Figure 3.**
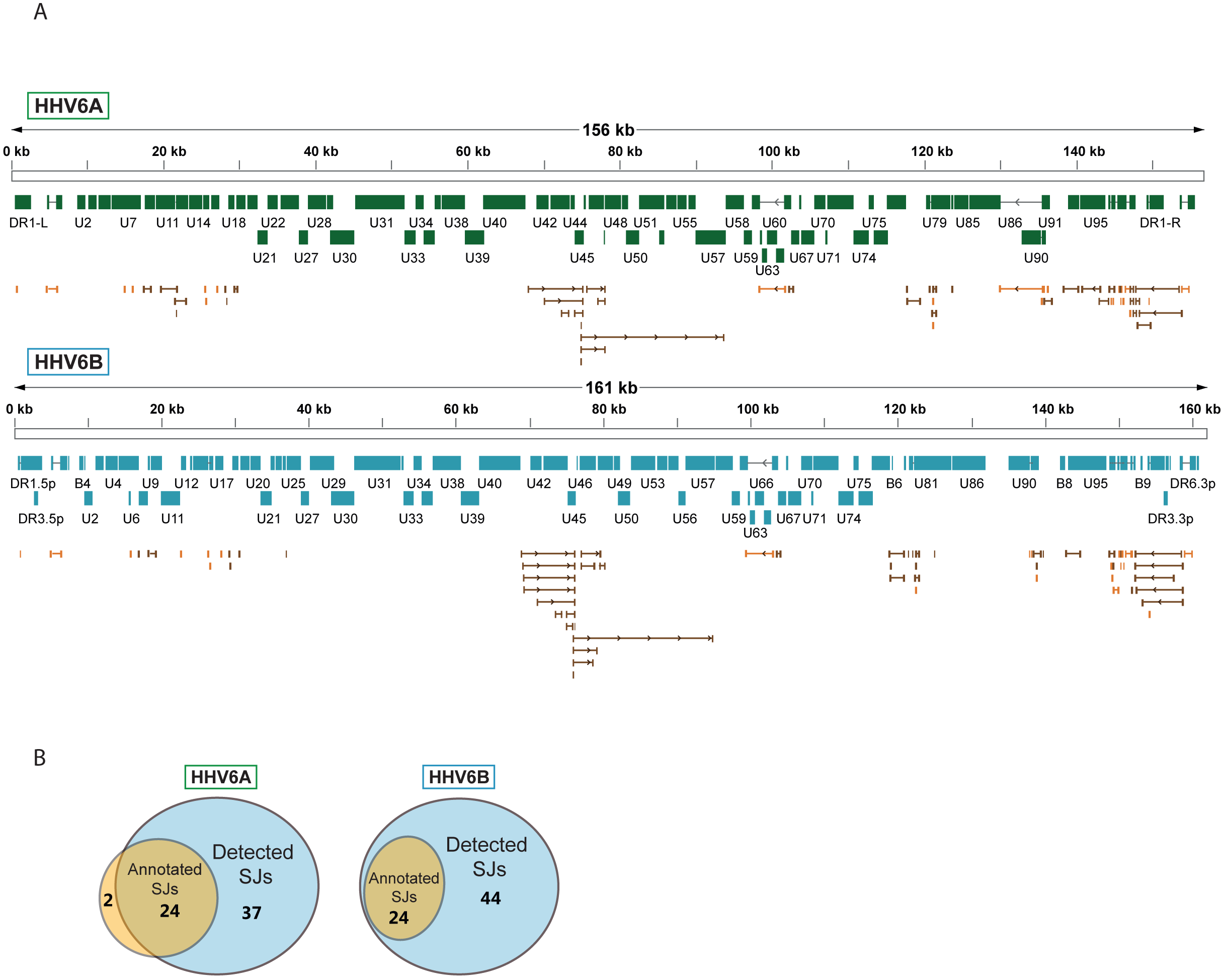
Splicing is abundant in HHV-6A and HHV-6B. (A) Splice junctions mapped using RNA-seq reads are shown throughout the genomes of HHV-6A and HHV-6B. Previously annotated splice junctions are marked in orange and novel splice junctions are marked in brown. (B) Diagrams displaying the numbers of previously annotated and detected splice junctions for HHV-6A and HHV-6B.

### Previously unrecognized HHV-6 encoded long non-coding RNAs (lncRNAs)

By examining the RNA-seq data, we discovered three highly expressed novel transcripts, that lack both observed or potential long ORFs, suggesting that these are likely lncRNAs. These three lncRNAs are conserved between HHV-6A and HHV-6B, and they all contain efficiently translated short ORFs (Figure 4A-C). The short length of these ORFs implies that the RNAs themselves probably constitute functional elements. One lncRNA, designated here as lncRNA1, initiates within HHV-6 origin of replication (Figure 4A) and therefore resembles in synteny to HCMV encoded lncRNA, RNA4.9 although it is much shorter (Figure S2A). This transcript is the most highly expressed RNA in both HHV-6A and HHV-6B (Figure S3), and its encoded short ORF contains the highest ribosome densities in the viral genomes (Table S3). The second lncRNA we identified, named here lncRNA2, is a spliced transcript that partially overlaps U18 (Figure 4B). The third lncRNA, designated lncRNA3, is transcribed between U77 and U79 (Figure 4C).

**Figure 4.**
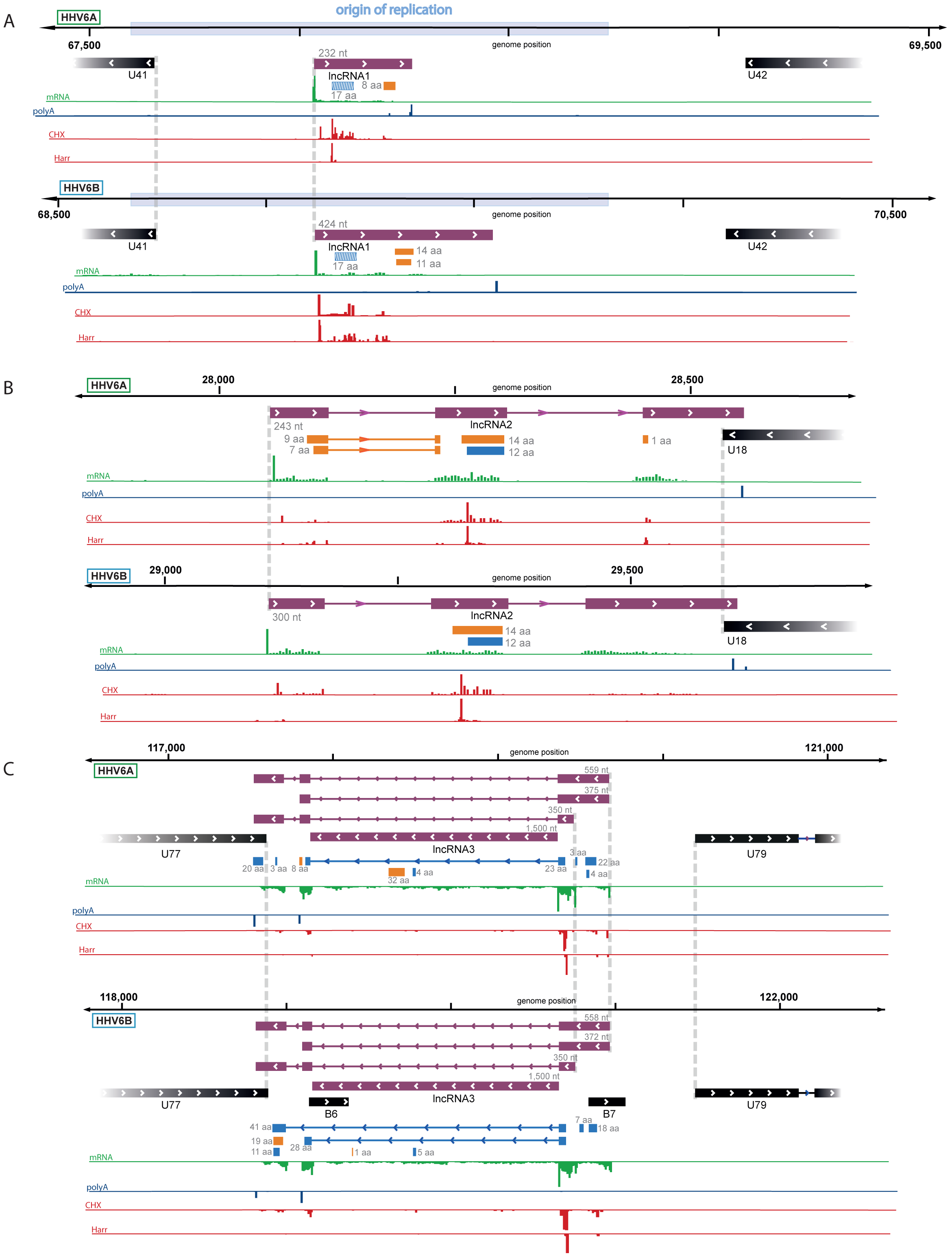
Identification of three highly abundant and conserved viral long non-coding RNAs (lncRNAs) Viral transcripts that appear to be lncRNAs are shown as purple rectangles. Reads from RNA-seq are presented in green and reads containing polyA are presented in blue. The ribosome profiling (CHX), and Harringtonine (Harr) profiles are presented in red. (A) A transcript initiating within the origin of replication. One putative ORF not detected by machine learning tools (see figure 5) is shown as a striped blue rectangle. (B) A spliced transcript initiating between U17 and U18. (C) Three possible isoforms of a spliced transcript with alternative splicing, initiation and termination, as well as a putative stable intron.

This lncRNA has multiple possible isoforms generated by two alternative TSSs, two alternative polyadenylation sites, and alternative splicing. Initial inspection of the RNA-seq data suggested that the intron is not efficiently spliced (Figure 4C). However by synteny, this lncRNA is homologous to HCMV encoded lncRNA5.0 and the Murine CMV encoded lncRNA7.2 (Figure S2B), shown to generate stable intronic RNAs which are not polyadenylated (Kulesza and Shenk, 2004, 2006).

Since our RNA-seq libraries were based on poly-A selection and therefore non-polyadenylated RNA molecules are under-represented, we suspected similar intronic RNA product might be generated from lncRNA3. To explore this possibility, we quantified the number of reads that span the exon-intron junction relative to the number of intronic reads, and found that in both HHV-6A and HHV-6B they comprise less than 10% of what is expected from retained intron isoforms (Figure 5A). Therefore, these intronic reads do not seem to originate from intron retention and rather indicate that lncRNA3 also generates a stable non-polyadenylated intron. To further explore this possibility, we extracted RNA from cells infected with HHV-6A or HHV-6B and measured the abundance of lncRNA3 intron in cDNA synthesized with random hexamers compared to cDNA synthesized with poly(dT) oligomers. Similar to the non-polyadenylated 18S ribosomal RNA, the intron RNA was detected at significantly higher levels in cDNA that was synthesized using random hexamers, while the polyadenylated lncRNA2 was more abundant or unchanged when poly(dT) oligomers were used in HHV-6A and HHV-6B, respectively (Figure 5B).

**Figure 5.**
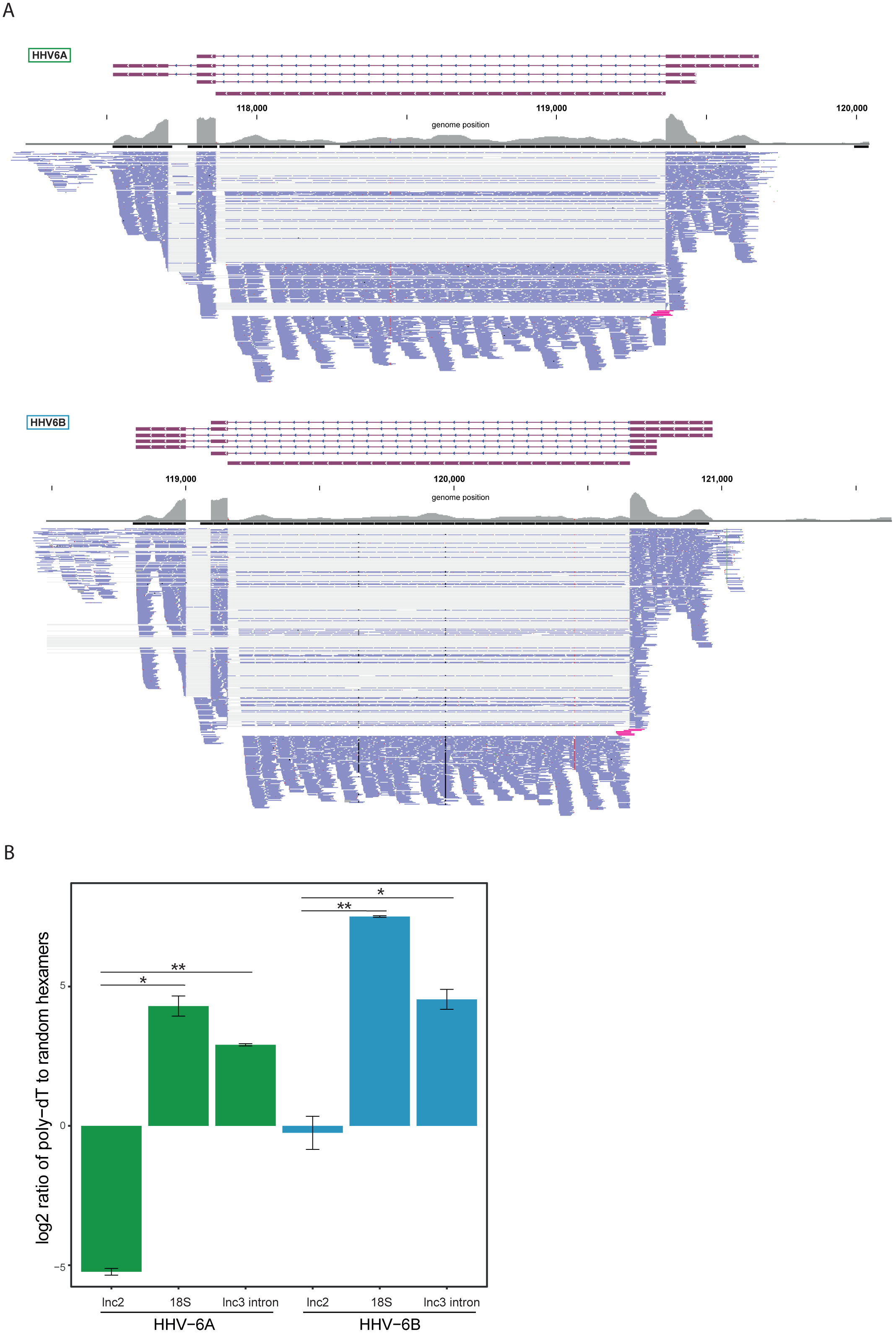
lncRNA3 generates a stable non poly adenylated intron. (A) RNA-seq reads aligned to the negative strand of lncRNA3 locus in both HHV-6A and HHV-6B are presented. Thin grey lines represent spliced reads, blue lines represent reads aligned to either the exons or intron, pink lines represent reads that span the first exon intron junction. In regions with very high coverage (>100 reads per 50 nt region) reads were downsampled so that maximum 100 reads per region are displayed. Grey bars represent the total reads coverage without omissions. (B) RT-qPCR measurements of the HHV-6A and HHV-6B lncRNA3 intron RNA. Values were normalized to the HHV-6 U21 gene. cDNA was prepared with either oligo-dT or random hexamers primers and the ratio of these measurements is presented. Error bars represent standard error of biological duplicates. p-values were calculated using Student’s t-test. * p-value < 0.05 and ** p-value < 0.01.

Taken together, our results show that HHV-6 viruses express three highly abundant lncRNAs, as was shown in other herpesviruses (Gatherer et al., 2011; Hutchinson and Tocci, 1986; Kulesza and Shenk, 2004; McDonough et al., 1985; Rawlinson and Barrell, 1993), and that one of these lncRNAs, lncRNA3, generates a stable non-polyadenylated intronic RNA that appears to be a conserved feature of betaherpesviruses.

### Systematic annotations of translated viral ORFs

To systematically define the full coding potential of HHV-6A and HHV-6B, we trained a support vector machine (SVM) model to identify translation initiation sites based on our Ribo-seq data sets, combining the actively translating ribosomes profile (CHX treatment), and initiation site enrichment (LTM and Harr) from cells infected with HHV-6 for 72 hours. The model was trained on a subset of the canonical viral ORFs that had high ribosome footprint coverage (see materials and methods). Using the trained SVM model, we predicted hundreds of translation initiation sites in each virus (Table S4 and S5). In these sites, we found strong enrichment of translation initiation at the canonical ATG start codon, as well as weaker but still significant enrichment for the near-cognate start codons (Figure 6A). Of the near-cognate start codons, CTG was the most common, similar to what was found in other herpesviruses (Arias et al., 2014; Stern-Ginossar et al., 2012; Whisnant et al., 2019) and in human cells (Fields et al., 2015). Of the previously annotated ORFs, we identified translation in 69 out of 88 HHV-6A ORFs and 63 out of 103 HHV-6B ORFs. The ORFs missing from the prediction were either reannotated, or hardly translated under the conditions we used (table S6). Since our detection is affected by the level of expression, it is likely these ORFs are expressed at low levels or translated under different conditions. In total, we identified 268 novel ORFs in HHV-6A and 216 novel ORFs in HHV-6B (Figure 6B). As expected, newly identified ORFs are shorter than the annotated ones (Figure 6C). Many of the novel ORFs we identified, were very short (<20 aa, 141 in HHV-6A and 111 in HHV-6B) and therefore are likely not functional at the polypeptide level. In addition, a large portion of the remaining ORFs are iORFs, translated within other ORFs (80 ORFs in HHV-6A and 67 in HHV-6B, figure 6B). Due to the nature of the ribosome movement on the RNA during active translation, the ribosome protected fragments of coding sequences display a three-nucleotide periodicity, with enrichment for reads aligned to the first base of each codon. The newly identified ORFs displayed similar periodicity to the previously annotated ORFs, which was not seen in RNA-seq reads, further validating that these ORFs likely represent bona-fide translation products (Figure 6D).

**Figure 6.**
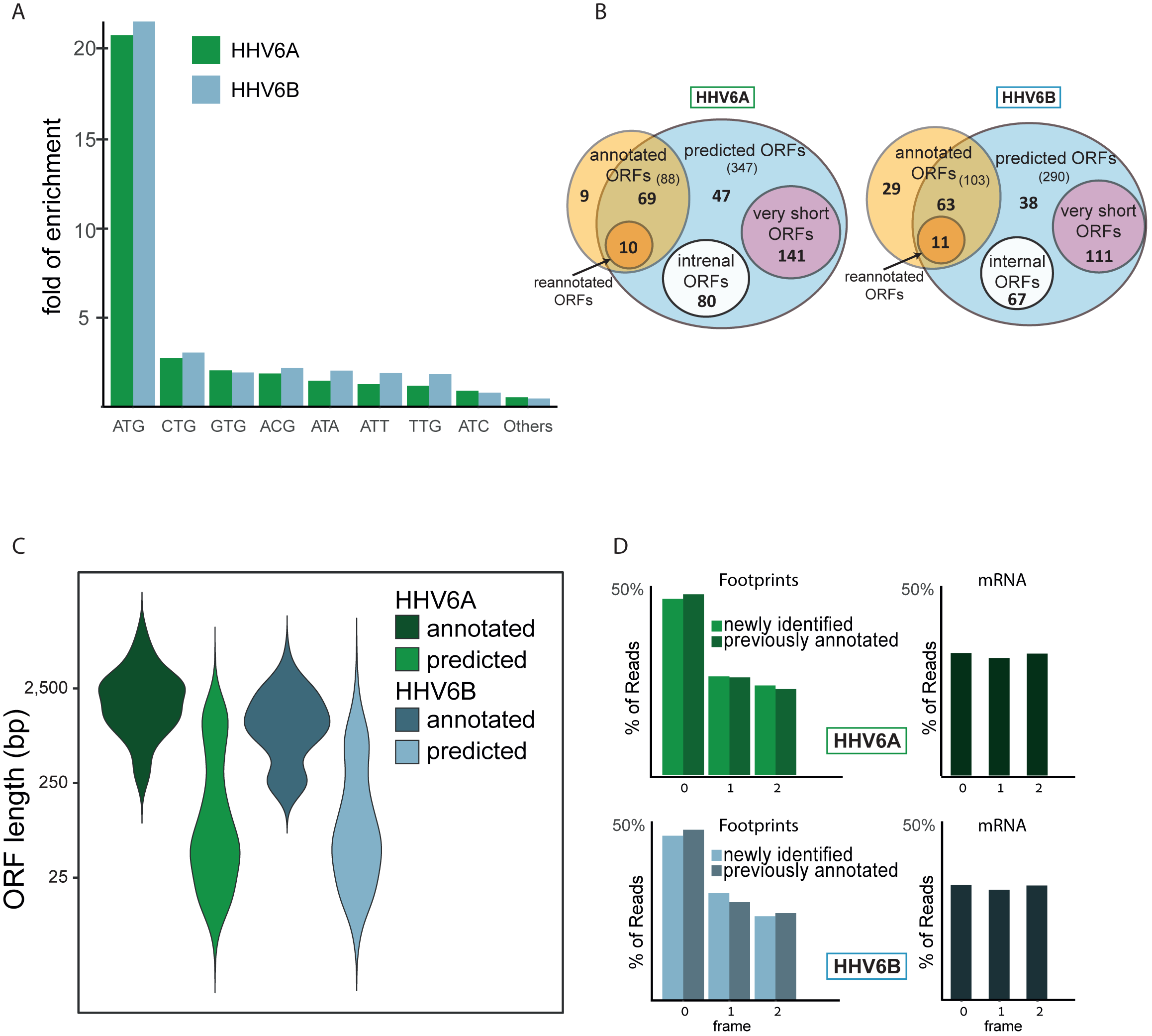
Identification of hundreds of novel HHV-6 ORFs. (A) Fold enrichment of AUG and near-cognate codons at predicted sites of translation initiation compared to their genomic distribution. (B) Venn diagrams summarizing the HHV-6 translated ORFs. (C) Size distribution of previously annotated ORFs (dark) and of newly identified ORFs (bright). (D) position of the ribosome footprint reads relative to the translated reading frame showing enrichment of the first position in the annotated ORFs (dark) as well as in the newly identified ones (bright). The mRNA reads were used as control and do not show enrichment.

### Pervasive use of alternative 5’ transcript ends controls viral gene expression

Gene expression during lytic herpesvirus infection is regulated in a temporal cascade. In order to explore the temporal kinetics of HHV-6 ORFs we performed a time course experiment and created Ribo-seq and RNA-seq libraries from HHV-6B strain Z29 infected Molt-3 cells at 5, 24 and 72 hours post infection (hpi). For these experiments we chose to focus on HHV-6B as it is more common and clinically relevant (Braun et al., 1997; Clark, 2016). The data in this experiment was highly correlated with our single time point experiment (Pearson’s R on log transformed data is 0.98 for RNA-seq 0.97 and for Ribo-seq, Figure S4).

Hierarchical clustering of viral coding regions by footprint densities along infection (a measure of the relative translation rates) revealed several distinct temporal expression patterns (Figure 7A and table S7). These temporal profiles largely agree with known kinetic classes (Yamanishi et al., 2013). Cluster 1 contains ORFs whose expression is relatively high at 5hpi compared to 24 and 72hpi, and this cluster includes most of the known immediate-early (IE) genes (U79, U90 and U95). A fourth IE gene, U85, was not efficiently translated at 5hpi and was assigned to cluster 2. Cluster 2 contains genes that are most highly expressed at 24hpi and is enriched in early genes. Clusters 3 and 4 contain genes that are mostly expressed at 72hpi and are both enriched in late genes; however, cluster 4 is composed of genes that are expressed almost exclusively at 72hpi. While most of the previously annotated late genes were assigned to these clusters, the DNA helicase/primase U43 and the large tegument protein U31 were previously annotated as late genes, but are shown here to reach peak translation at the early 24hpi timepoint.

**Figure 7.**
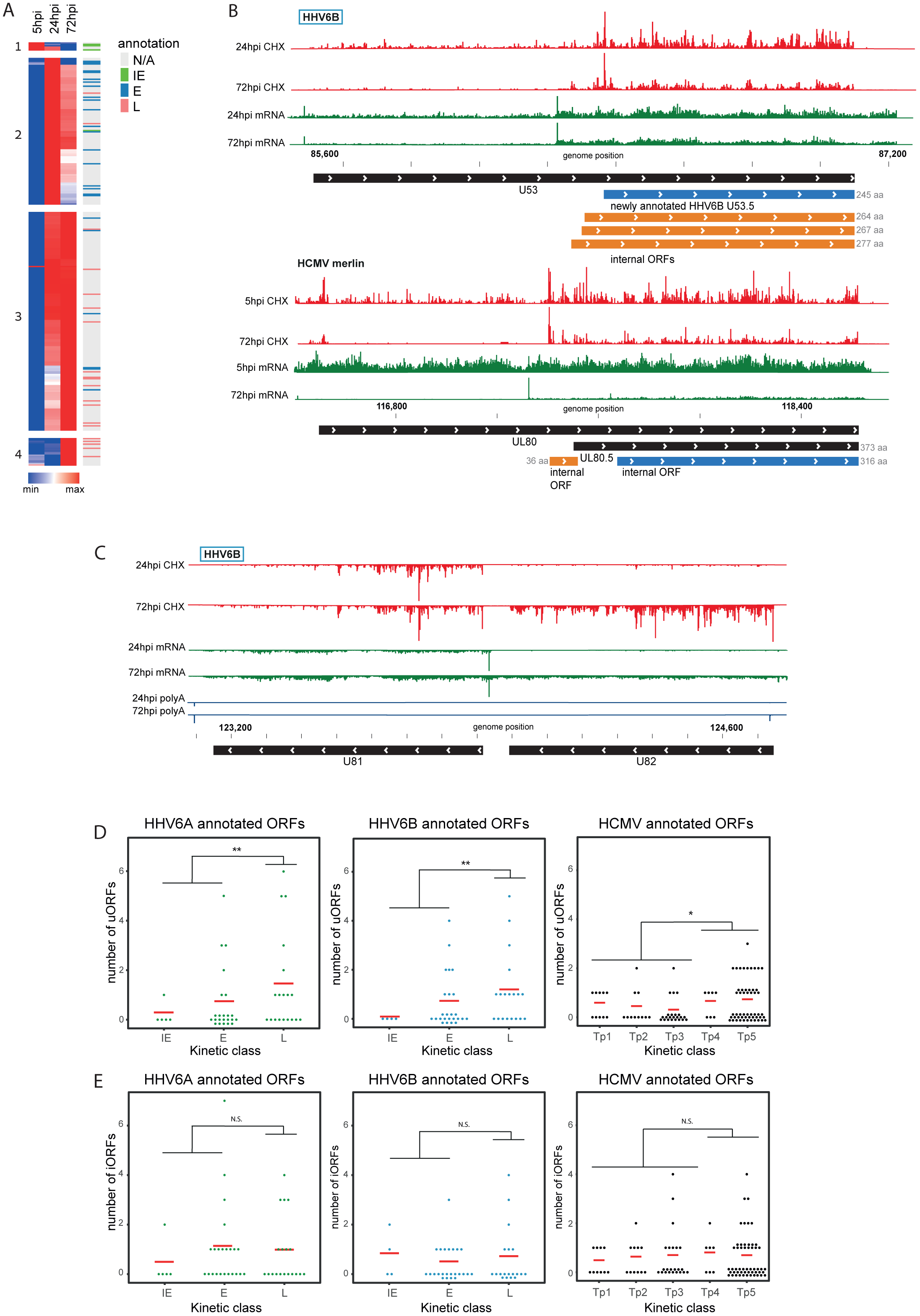
Temporal regulation of viral gene expression is driven by pervasive use of alternative 5’ ends. (A) Heatmap of ribosome occupancy of HHV-6B ORFs clustered by relative expression levels at 5, 24 and 72hpi. ORFs with known kinetic class were annotated on the right as immediate early (IE, green), early (E, blue), late (L, pink), or unknown (N/A, grey). The cluster number appears on the left. (B and C) The ribosome occupancy (red) and mRNA profiles (green) are shown (B) around U53 loci at different hours post infection (marked on the left) and around its HCMV homolog, UL80 (C) and around U81 and U82 loci. (D and E) Dot plots showing the number of uORFs (D) and iORFs (E) of each canonical viral ORF with known kinetic class for HHV-6A, HHV-6B and HCMV. P value was calculate using proportion test. * for p-value < 0.05, ** for p-value < 0.01 and N.S for non-significant.

We previously demonstrated that pervasive use of alternative 5’ ends in HCMV transcripts is critical for the tight temporal regulation of viral gene expression and production of alternate protein products (Stern-Ginossar et al., 2012). We observed similar phenomena in the temporal regulation of several HHV-6B genes. For example, the U53 gene contains newly identified iORFs, one of which initiates at an ATG, and is an orthologue of the annotated HHV-6A U53.5 ORF (Figure 7B). Relative to the main U53 ORF, these iORFs are translated more efficiently at 72hpi than at 24hpi. This could be explained by a temporal shift in the relative frequency of initiation at two TSSs, one of which is upstream of the U53 start codon from which the main U53 can be translated, and another downstream of the U53 start codon allowing translation of the iORFs but not of the main U53 ORF. Notably, we found the same pattern in the HCMV homolog, UL80 ((Stern-Ginossar et al., 2012), figure 7B). A similar form of regulation is seen in the HHV-6B locus coding for the U81 and U82 ORFs in which we found two TSSs. One TSS is immediately upstream of U81 creating an RNA that is mainly expressed at 24hpi, facilitating the translation of U81. At 72hpi a second TSS is also present, giving rise to translation of U82 (Figure 7C). Temporal regulation of 5’ ends was also found for the HHV-6B U51 and its uoORF, which is also conserved in its HCMV homolog UL78 ((Stern-Ginossar et al., 2012), Figure S5).

### uORFs are enriched in betaherpesvirus late genes

Among the newly identified ORFs many are iORFs and uORFs. Since the high abundance of these ORFs may be associated with the fidelity of the ribosome, or alternatively, can be related to specific translation regulation, we examined whether these translation events are enriched in specific kinetic classes. Each uORF and iORF was assigned to a canonical transcript; iORFs were assigned to the canonical ORF in which they reside, and uORFs where assigned to a canonical ORF if they were located upstream of its translation initiation (Table S8). For both HHV-6A and HHV-6B we found an enrichment of uORFs in the 5’UTRs of late genes compared to earlier kinetic classes (Figure 7D, p-value < 0.01, proportion test). In contrast, there was no enrichment for the presence of iORFs in any kinetic class (Figure 7E, p-value > 0.3) negating the option that the enrichment we found for uORFs is due to a bias in our approach or to general unspecific increase in translation initiation rate of ribosomes. We further extended this analysis to HCMV ORFs and found that uORFs but not iORFs are significantly enriched in 5’UTRs of late genes, similar to what we see in HHV-6 (Figure 7D and 7E). These results suggest a potential mechanism for translation regulation of late viral genes, utilizing uORFs, which is conserved among betaherpesviruses.

### The presence of iORFs and uORFs is conserved among betaherpesvirus genes

Using our comprehensive transcriptome and translatome data we uncovered hundreds of novel ORFs in HHV-6A and in HHV-6B. We next examined whether the presence of these ORFs is conserved between these two HHV-6 species. We found that the number of iORFs and uORFs in HHV-6A and HHV-6B homolog ORFs are well correlated, indicating a high level of conservation of these translation events between these two viruses (p < 10^-15^ for uORFs and p < 10^-10^ for iORFs, figure 8A). Several homolog ORFs have multiple conserved iORFs and/or uORFs (Fig 8B and figure S6). We also found some features that are conserved in HCMV. In five iORF-containing HHV-6 genes and in four uORF-containing HHV-6 genes, the HCMV homologs also contained similar iORFs or uORFs (Figure S7). One of the HHV-6/HCMV homolog ORF pairs containing a conserved uORF is U51 and its HCMV homolog UL78 (Figure 8C), which interestingly also show conserved kinetics along infection suggesting a potential regulatory mechanism conserved between these viruses (Figure S5). Altogether, the conserved presence of several uORFs and iORFs suggests that their occurrence is not random, and it is likely that these represent a functional module that plays a role in regulating herpesvirus protein expression.

**Figure 8.**
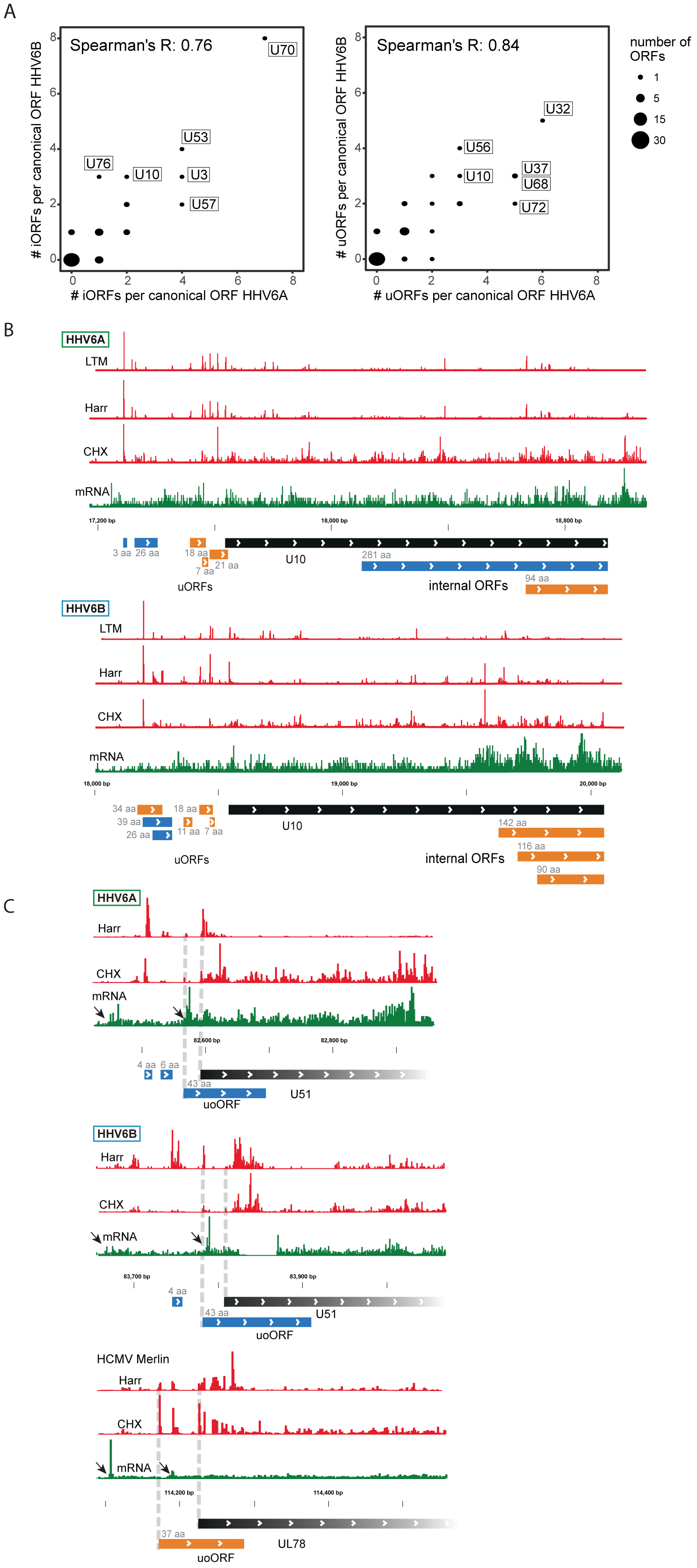
Numerous iORFs and uORFs are conserved between betaherpesviruses. (A) Correlation between the number of iORFs and uORFs of canonical ORFs in HHV-6A and HHV-6B (55 shared canonical ORFs in total). Dot size indicates the number of canonical ORFs with the indicated number of iORFs or uORFs in the two viruses. (B-C) Selected examples of novel internal or upstream initiation events that are conserved between HHV-6A and HHV-6B are presented. Shown in black rectangles are canonical ORFs, in blue are novel ORFs initiating at an AUG codon, and in orange are novel ORFs initiating at a near-cognate start codon. ORF sizes are written in grey. The ribosome occupancy (CHX), harringtonine (Harr), LTM and mRNA profiles are shown in red and the mRNA profile is shown in green (B) at U10 locus for both HHV-6A and HHV-6B and (C) at the U51 locus in HHV-6A and HHV-6B and its HCMV homolog U78. The gap in RNA reads in HHV-6B U51 is due to a base insertion relative to the reference, preventing read alignment to the region.

## Discussion

Decoding the transcriptional and translational landscape of any virus is a fundamental step in studying its biology and pathogenesis. For many herpesvirus genomes, traditional annotations have relied on the identification of canonical translational start codons and arbitrary size restriction to define viral open reading frames (ORFs). Laborious follow-up molecular work revealed the transcriptional architecture of individual genomic loci, but for most HHV-6 genes annotations are still based on these initial *in-silico* ORF predictions. In recent years, major advances in high-throughput sequencing approaches have revealed that the transcriptome and translatome of herpesviruses are extremely complex, encompassing large numbers of overlapping transcripts, extensive splicing and many non-canonical translation products (Arias et al., 2014; Balázs et al., 2017; Bencun et al., 2018; Depledge et al., 2019; Gatherer et al., 2011; Kara et al., 2019; O’Grady et al., 2019, 2016; Tombácz et al., 2017; Whisnant et al., 2019). Our own work in which we employed ribosome profiling and systematic transcript analysis to decipher HCMV genome complexity revealed a rich collection of coding sequences that include many viral short ORFs (sORFs), uORFs and alternative translation products that generate extensions or truncations of canonical proteins (Stern-Ginossar et al., 2012).

Here, using RNA-Seq and ribosome profiling measurements along HHV-6 infection, we provide a comprehensive map of HHV-6A and HHV-6B coding elements over the lytic life cycle. In agreement with the complexity of other herpesviruses that have been analyzed using ribosome profiling approaches, we identified 268 and 216 novel viral ORFs that are expressed during HHV-6A and HHV-6B lytic infection, correspondingly. Furthermore, our transcriptome analyses enabled mapping of the full landscape of HHV-6 splice junctions and the identification of three virally encoded lncRNAs. Our data further show that in similarity to our findings in HCMV (Stern-Ginossar et al., 2012), the pervasive use alternative 5’ ends plays a major role in HHV-6 genomes in production of distinct polypeptides from single genomic loci. Like alternative splicing, this mechanism can expand protein diversity and contribute to virus complexity by allowing multiple distinct polypeptides to be generated from a single genomic locus. Overall, our revised experimental annotations will facilitate functional studies on HHV-6 ORFs and transcripts as well as their regulation.

This wealth of novel elements requires a more precise dissection of the components that are likely to be functional. The issue of functional relevance still represents a major challenge in these systematic experimental annotations. Our analysis of three betaherpesviruses allowed us, for the first time, to highlight some conserved features that may point towards functional importance.

A large portion of the novel translated ORFs we identified are uORFs. uORFs are widely recognized as *cis*-regulatory elements and their presence generally correlates with reduced translation of the primary ORF, but there are instances in which they associate with increased translation (Young and Wek, 2016). Despite their pervasiveness, only a few viral uORFs have been studied in detail (Geballe et al., 1986; Kronstad et al., 2013). We show that genes that contain uORFs and the number of uORFs are largely conserved between HHV-6A and HHV-6B. In addition, we reveal that both in HHV-6 and in HCMV, uORFs appear to be especially abundant in late viral genes. The surplus of uORFs and their preferred use specifically at late time points of infection indicate that many of them may have a functional role in controlling viral gene expression, probably when cellular stress pathways are engaged at late time points of infection. Given the overall high representation of sORFs, iORFs and uORFs in the viral genome, which are often translated from near canonical start codons, it is tempting to speculate that the extensive use of non-canonical start codons, particularly late during infection, is caused by changes in translation permissiveness.

We identified three conserved HHV-6 encoded lncRNAs, signifying lncRNAs are probably a shared feature of all herpesviruses (Tycowski et al., 2015). lncRNAs are still an enigmatic group of RNA molecules that do not form a well-defined class of genes, and mechanistically most lncRNAs, including viral lncRNAs, remain poorly characterized. Unlike mammalian lncRNAs, that as a group are significantly less abundant than canonical mRNAs (Mukherjee et al., 2017), in herpesviruses lncRNAs represent the most abundant group of viral transcripts (Gatherer et al., 2011; Tycowski et al., 2015). These high expression levels allude to essential roles for virally encoded lncRNAs during infection. The three HHV-6 encoded lncRNAs we identified are highly expressed and present distinct features; lncRNA1 is relatively short (232bp in HHV-6A and 424bp in HHV-6B) and unspliced, lncRNA2 is composed of three exons that are efficiently spliced, and lncRNA3 represents a complex locus with two different TSSs and polyadenylation sites, three alternatively spliced exons and a stable intron. Interestingly, by synteny lncRNA1 and lncRNA3 seem related to HCMV encoded lncRNAs. lncRNA1 resembles the HCMV RNA4.9 as both are transcribed from the viral origin of replication at the same orientation. This similarity implies a possible conserved role of lncRNA transcription in betaherpesviruses origin of replication although they are very different in length (RNA4.9 is 4.9KB long). lncRNA3 is an orthologue of HCMV encoded RNA5.0 both in synteny and in the production of a stable intron. RNA5.0 has previously been shown to generate a stable intron that is not required for HCMV replication in fibroblasts (Kulesza and Shenk, 2004). A murine cytomegalovirus 7.2-kb ortholog of RNA5.0 was identified which also generates a stable intronic RNA. Mutant MCMV viruses lacking this stable intron RNA replicated normally in culture, but exhibited defect in establishing a persistent infection in-vivo (Kulesza and Shenk, 2006). Our results indicate that the production of a stable intronic RNA from this locus is a conserved feature of betaherpesviruses, implying a central function. Importantly, the notion that this non-coding region is conserved in betaherpesviruses and therefore likely represents a functional component was already specified 15 years ago (Dolan et al., 2004). Although we used poly-A selection and these intronic RNAs are not polyadenylated, the oligo(dT) beads may have bound to homopolymeric stretches of A residues found throughout the intron sequence. However, it is likely that our RNA-seq measurement underestimate the true abundance of theses intronic RNAs and it is probable these are the most abundant transcripts in infected cells. This high abundance together with the wide conservation in betaherpesviruses make the molecular and functional characterization of these viral intronic RNAs highly interesting. There is little known about the mechanisms by which stable intronic RNAs may operate (Osman et al., 2016) but one possibility is that these RNAs sequester spliceosomes or specific splicing components that cause changes in the cellular splicing activity.

In conclusion, we provide a comprehensive annotation of HHV-6 transcripts and ORFs and highlight conserved translation patterns and non-coding RNAs that may have central shared functions in all betaherpesviruses.

## Acknowledgements

We thank the members of the Stern-Ginossar lab for critical reading of the manuscript. This research was supported by the ICORE (Chromatin and RNA Gene Regulation, N.S-G.) and the Israeli Science Foundation (1526/18, N.S-G.). N.S-G is incumbent of the Skirball career development chair in new scientists.

## Author contributions

Y.F, D.S, J.T-S, M.S, O.M and N.S-G. conceived experiments and interpreted data. D.S and J.T-S performed ribosome profiling experiments. Y.F and A.N analyzed the data. Y.F and N.S.G. wrote the manuscript with contribution from all other authors.

## Materials and methods

### Cell lines and virus strains

HSB-2 Cells from Electro-Nucleonics, Inc. (Barre-Sinoussi et al., 1983) and Molt-3 cells (ATCC CRL1552) were maintained at 37°C in 5% (vol/vol) CO2, in RPMI 1650 medium (Biological Industries) supplemented with 10% heat-inactivated fetal bovine serum (Life Technologies), 2mM L-glutamine (Biological Industries), 1mM sodium pyruvate, 0.1mg/mL streptomycin and 100 U/mL penicillin (Biological Industries).

HHV-6A strain GS and HHV-6B strain Z29 were maintained in HSB2 and Molt-3 cells, respectively. For viral propagation, infected cells were added to uninfected cells at a ratio of 1:10 every 3 or 4 days. All viruses and cell lines were obtained from the NIH AIDS reagent program, Division of AIDS, NIAID, NIH.

### Preparation of Ribosome profiling and RNA sequencing samples

Samples were prepared by co-incubating either HSB-2 or Molt-3 at a density of 1-1.5M cells per mL with cells infected with HHV-6A and HHV-6B, respectively, at a 1:5 ratio, for 72 hours.

For RNA-seq, cells were harvested with Tri-Reagent (Sigma-Aldrich), and total RNA was extracted. For Ribo-seq libraries, cells were treated with either 50µM LTM for 30 minutes or 2µg/mL Harringtonine for 5 minutes, for the translation initiation libraries (LTM and Harr), or left untreated for the translation elongation libraries (CHX). All three samples were subsequently treated with 100µg/mL cycloheximide for 1 minute. Cells were placed on ice immediately after treatment, centrifuged and washed twice with PBS containing 100µg/mL cycloheximide. Subsequent lysis, Ribo-seq and RNA-seq library generation were performed as previously described (Ingolia et al., 2011).

For HHV-6B infection kinetics, virus containing supernatant was collected from Molt-3 cells infected for four days. Six samples of 250,000 Molt-3 cells were incubated in 50µL of the viral supernatant each, for 30 minutes at 4°C and then for 45 minutes at 37°C. After infection, the cells were incubated in RPMI at a cell density of 1 million per 1.5 mL. Cells were harvested at 5, 24 and 72 hours post infection, and CHX and RNA-seq libraries were generated as described above. Prepared libraries were sequenced on the illumina NextSeq 550 with at least 61 nt single-end reads.

### Sequence alignment, normalization, metagene analysis and clustering and visualization

Sequencing reads were aligned as previously described (Tirosh et al., 2015). Briefly, linker and poly-A sequences were removed and the remaining reads were aligned to the human and the viral reference genomes (HHV-6A KC465951.1, HHV-6B AF157706.1) using Bowtie v1.1.2 (Langmead et al., 2009) with maximum two mismatches per read. Reads that were not aligned to the genome were aligned to the transcriptome, taking into account all the new identified splice junctions. Reads aligned to multiple locations were ignored. Sequencing data was visualized using IGV integrative genomics viewer (Robinson et al., 2011).

For the metagene analysis only genes with more than 100 reads were used. Each gene was normalized to its maximum signal and each position was normalized to the number of genes contributing to the position.

For the time-course clustering, footprints counts were normalized to units of reads per million (RPM) in order to normalize for sequencing depth. To avoid noise arising from low viral gene expression at 5hpi, ORFs with less than six reads at this time point were considered to have zero reads. Morpheus (https://software.broadinstitute.org/morpheus) was used to perform hierarchical gene clustering with one minus Pearson correlation as metric and complete linkage method.

For comparing transcript expression level, mRNA and footprint counts were normalized to units of reads per kilobase per million (RPKM) in order to normalize for gene length and for sequencing depth.

Single nucleotide mutations in RNA-seq were identified (Mizrahi et al., 2018) and positions with at least 10 reads that had a different base than the reference in 95% or more of the reads are listed in tables S9 and S10. Lists of deletions and insertions that scored 20 or above in the TopHat output are also in tables S9 and S10.

### Identification of splice junctions

RNA-seq results were analyzed using TopHat v2.1.1 (Kim et al., 2013; Trapnell et al., 2009) with no coverage search, a minimum intron size of 15bp, and STAR v2.5.3a (Dobin et al., 2013) with default parameters. Splice junctions were chosen for the final annotations if they score 20 or higher in both STAR and TopHat, and if the intron length was less than 3.5Kb (to filter out artificial splice junctions between the viral repeat regions). We also included splice junctions that were detected and were previously known but did not pass the threshold (five junctions in HHV-6A and five in HHV-6B). Two additional previously annotated HHV-6B splice junctions that were not detected were added to the final list.

### Prediction of translation initiation sites

Translation initiation sites were predicted as previously described (Ingolia et al., 2011; Stern-Ginossar et al., 2012). Briefly, a support vector machine model was trained to identify initiation sites based on normalized footprint profiles of the CHX, Harr and LTM samples. A positive example set was composed of previously annotated translation initiation sites that were also well translated in our data (at least seven read counts in the normalized Harr peak, 39/58 ORFs for HHV-6A and 31/47 for HHV-6B). 10 negative examples were computed for each positive example. 2/3 of the combined set of positive and negative examples was used as a training set for the prediction model, using a radial basis kernel, γ = 2, C = 50, relative positive example weighing of 1.0, and without iterative removal and retraining. Initiation sites that scored less than 0.5 were discarded. The remaining 1/3 of the example set was used for cross-validation, which showed 37% and 25% false negative rate, and 2% and 5% false positive rate for HHV-6A and HHV-6B respectively. The trained classifier was then applied to all plus and minus strand codons that had at least 7 normalized Harr read counts. ORFs were then defined by extending each initiating codon to the next in-frame stop codon, and incorporating any intervening splice junctions. Previously annotated ORFs that were not recognized by the trained model but presented observable translation in manual inspection were added to the final ORF list (table S11).

### Comparison of uORF and iORF conservation and kinetics

uORFs were curated by selecting all ORFs of the predicted ORFs initiating in the 200bp region upstream of each previously annotated ORF that are shorter than 200bp. iORFs were curated by selecting ORFs longer than 20aa initiating within each previously annotated ORF. The total number of iORFs and uORFs for each main ORF was summed.

The comparison of HHV-6 annotations to HCMV was based on previously published Ribo-seq, RNA-seq and annotations of HCMV merlin strain (Stern-Ginossar et al., 2012; Tirosh et al., 2015). Text-book published lists were used to identify HHV-6 and HCMV homolog ORFs, as well as to determine the kinetic classes for previously annotated HHV-6 ORFs (Yamanishi et al., 2013). HCMV kinetic class annotations were taken from proteomics based publication (Weekes et al., 2014). Data for murine CMV lncRNA7.2 expression is from Tai-Schmiedel et al. unpublished.

All plot and statistical tests were done using R 3.6.0 (R Core Team, 2019; Wickham, 2016) on Rstudio 1.0.143 (RStudio Team, 2015).

### Real-time PCR

Total RNA was isolated from 72hpi Infected cells using Tri-Reagent (Sigma-Aldrich). Reverse transcription was performed with qScript Flex cDNA kit (Quantabio), using either oligo-dT or random primers, as described for each sample. Real time PCR was performed using the SYBR Green master-mix (ABI) on a real-time PCR system StepOnePlus (life technologies), with the following primers:

lncRNA3-6A F AAAAGGACAAGAGCAGCCGC

lncRNA3-6A R ACTCGTATCACCTACCTCTCTCTAC

lncRNA3-6A F GGTATCGGGGTAAGAATAAGATGACG

lncRNA3-6A R AAAAGGACAAGAGCAGCCGC

lncRNA2-6B F CAAAACGGTCTCACTGCTCC

lncRNA2-6B R TCTATAAAGTGCCGTGAGTGC

lncRNA2-6A F CGACAAAACAAAATAGTCCCACT

lncRNA2-6A R ATGGAAAAGGTGGTCGTGGA

U21-6B F CCGCACCCATGAACATAAGG

U21-6B R ATGATGTGACGTGGGGACTT

U21-6A F CCAGCCACCTAGAGAACGAA

U21-6A R TTGGGCTGAACTCTCGACAT

18S F CTCAACACGGGAAACCTCAC

18S R CGCTCCACCAACTAAGAACG

Results in CT were normalized to the U21 for sample virus and to oligo-dT cDNA for each replicate.

### Data availability

All next-generation sequencing data files were deposited in Gene Expression Omnibus under accession number GSE135363

---

**Figure S1. Novel splice junctions result in reannotation of HHV-6 ORFs**

Ribo-seq reads are shown in red and RNA-seq reads are shown in green for (A) the U7-9 locus of HHV-6A and HHV-6B. (B) the U12-13 locus of HHV-6A. Black rectangles mark canonical annotations of open reading frames, blue rectangles mark novel ORFs initiating at an AUG codon, and orange rectangles mark novel ORF initiating at a near-cognate start codon. Transcripts are shown in grey. RNA-seq read alignments in BAM format are shown at the bottom, thin grey lines represent spliced reads, pink lines are reads aligned to the positive strand and blue lines are reads aligned to the negative strand.

**Figure S2. Conservation by synteny of newly discovered HHV-6 lncRNAs**

(A and B) Reads from RNA-seq are presented in green. Black rectangles mark canonical ORFs and purple rectangles mark the putative lncRNAs (A) a lncRNA initiating within the lytic origin of replication of HHV-6A, HHV-6B and HCMV (RNA4.9). (B) a spliced lncRNA, likely generating a stable intron, transcribed from the locus between the viral helicase gene and a conserved early phosphoprotein gene in HHV-6A, HHV-6B, HCMV and Murine CMV (MCMV).

**Figure S3. RNA abundance of canonical ORFs and viral lncRNAs is conserved between HHV-6A and HHV-6B**

Scatter plot of normalized RNA expression levels of canonical HHV-6 ORFs and novel lncRNAs. Grey dots represent ORFs, colored dots represent lncRNAs (lncRNA1 in red, lncRNA2 in green and lncRNA3 in blue).

**Figure S4. RNA abundance and ribosome footprint coverage correlate well between replicates.**

Scatter plot of RNA-seq and CHX Ribo-seq reads of canonical HHV-6 ORFs and novel lncRNAs. Grey dots represent ORFs, colored dots represent lncRNAs (lncRNA1 in red, lncRNA2 in green and lncRNA3 in blue).

**Figure S5. Conserved temporal regulation of translation from uoORF**

The ribosome occupancy and mRNA profiles are shown around the HHV-6B U51 locus and the UL78 HCMV locus, at different infection times (marked on the left). CHX Ribo-seq reads are presented in red and RNA-seq reads are presented in green. Black rectangles represent canonical annotations, blue rectangles represent novel ORF initiating at an AUG codon and in orange rectangles represent ORFs initiating at a near-cognate start codon.

**Figure S6. Viral loci with conserved presence of multiple uORFs and iORFs**

Ribo-seq reads (red) and RNA-seq (green) of several virus loci. Black rectangles represent canonical annotations, blue rectangles represent novel ORF initiating at an AUG codon and in orange rectangles represent ORFs initiating at a near-cognate start codon. ORF sizes are written in grey. (A) multiple in-frame iORFs within U70 in HHV-6A and HHV-6B. (B) multiple uORFs upstream of U32 ORF in HHV-6A and HHV-6B.

**Figure S7. Synteny conservation of uORFs and iORFs between HHV-6 and HCMV**

Correlation between the number of iORFs and uORFs of canonical HCMV and HHV-6 ORFs (26 canonical main ORFs in total). Dot size indicates the number of canonical ORFs with the indicated number of iORFs or uORFs in the two viruses. (A) HHV-6A and HCMV uORFs, (B) HHV-6B and HCMV uORFs, (C) HHV-6A and HCMV iORFs, and (D) HHV-6B and HCMV iORFs.

## Supplementary files

### File S1 and S2: updated ORF annotations

Bed format file of genomic loci of ORFs in the genomes of HHV-6A (S1) and HHV-6B (S2) curated using SVM model predictions with manual modifications, see materials and methods.

### File S3 and S4: lncRNA annotations

Bed format file of genomic loci of newly identified lncRNAs in the genomes of HHV-6A (S3) and HHV-6B (S4).

