## Supplementary figures and images for "Comprehensive Annotations of Human Herpesvirus 6A and 6B Genomes Reveal Novel and Conserved Genomic Features"

### Supplemental figures

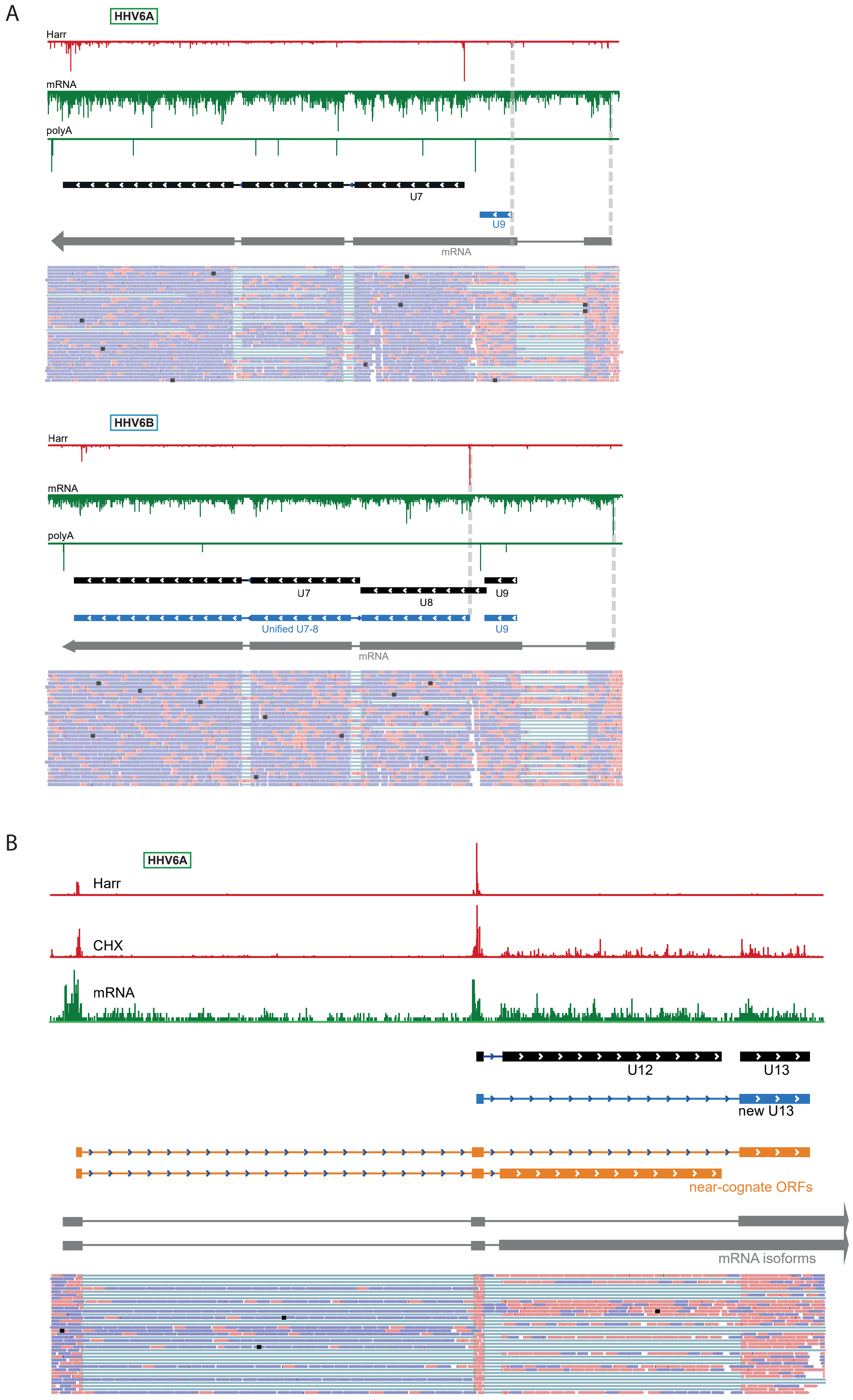

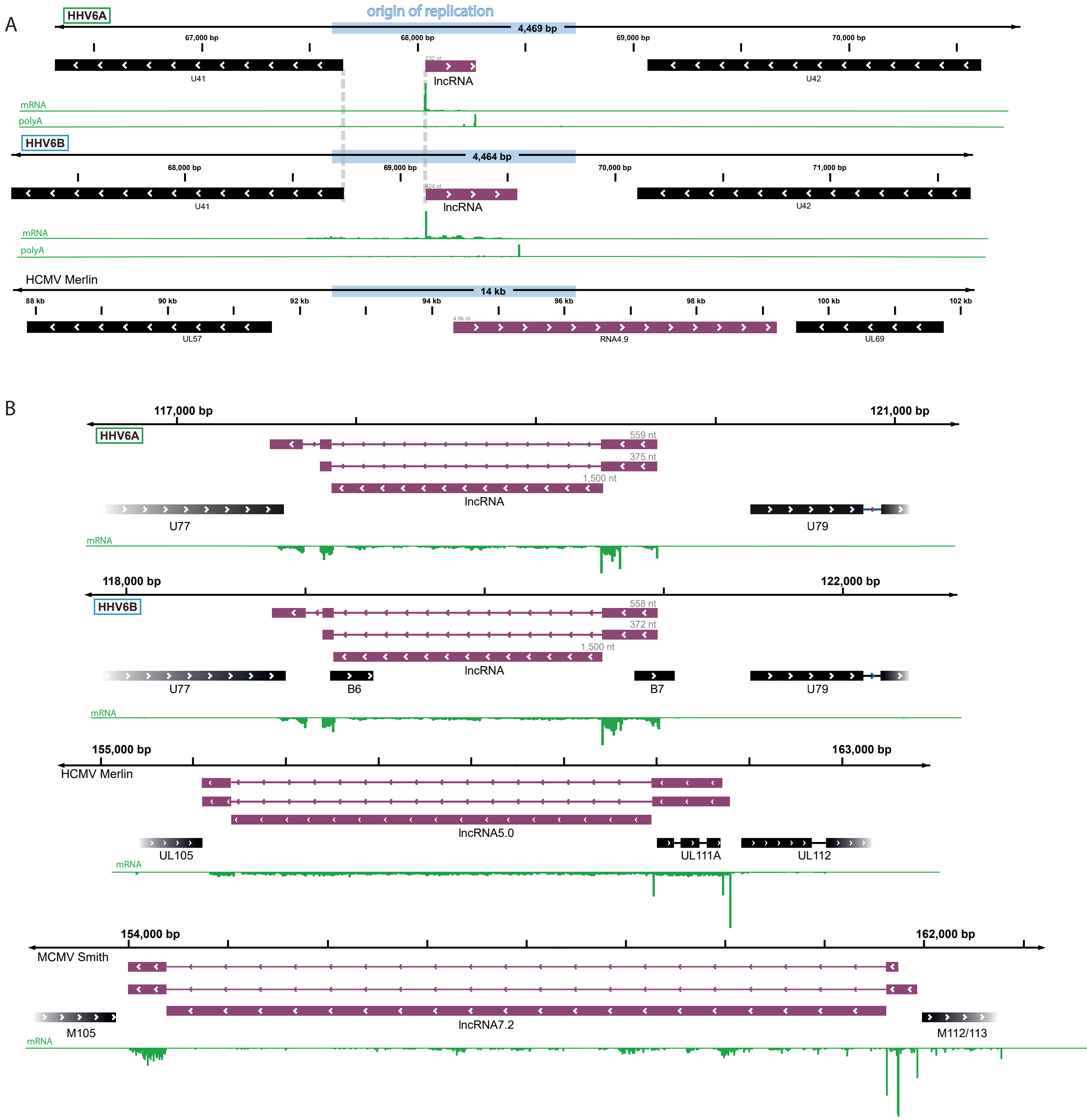

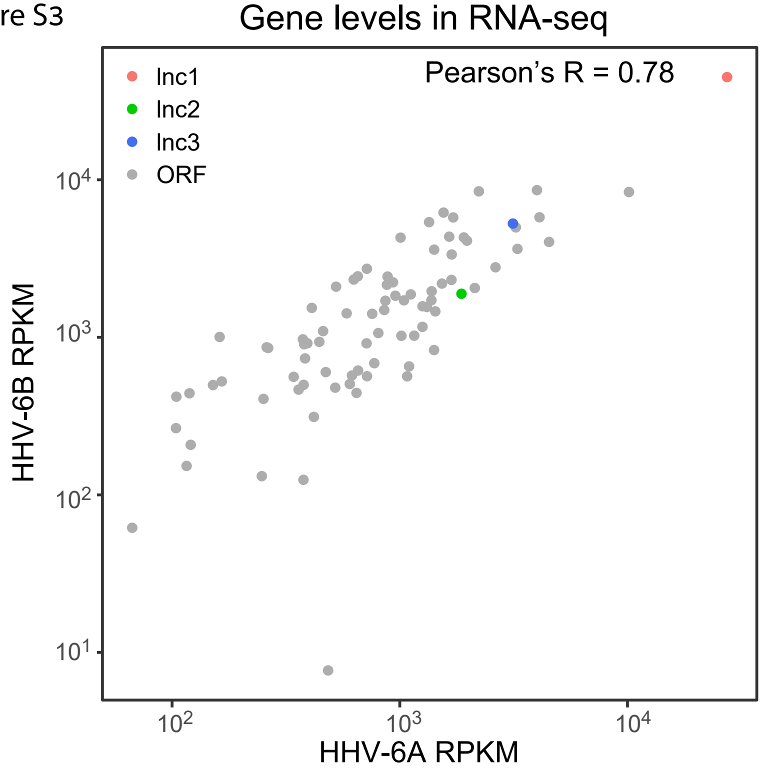

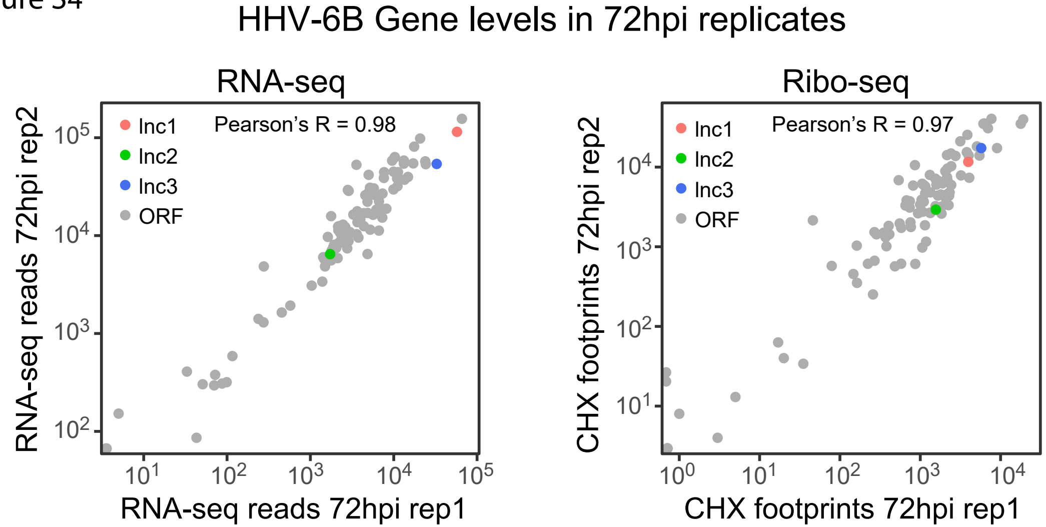

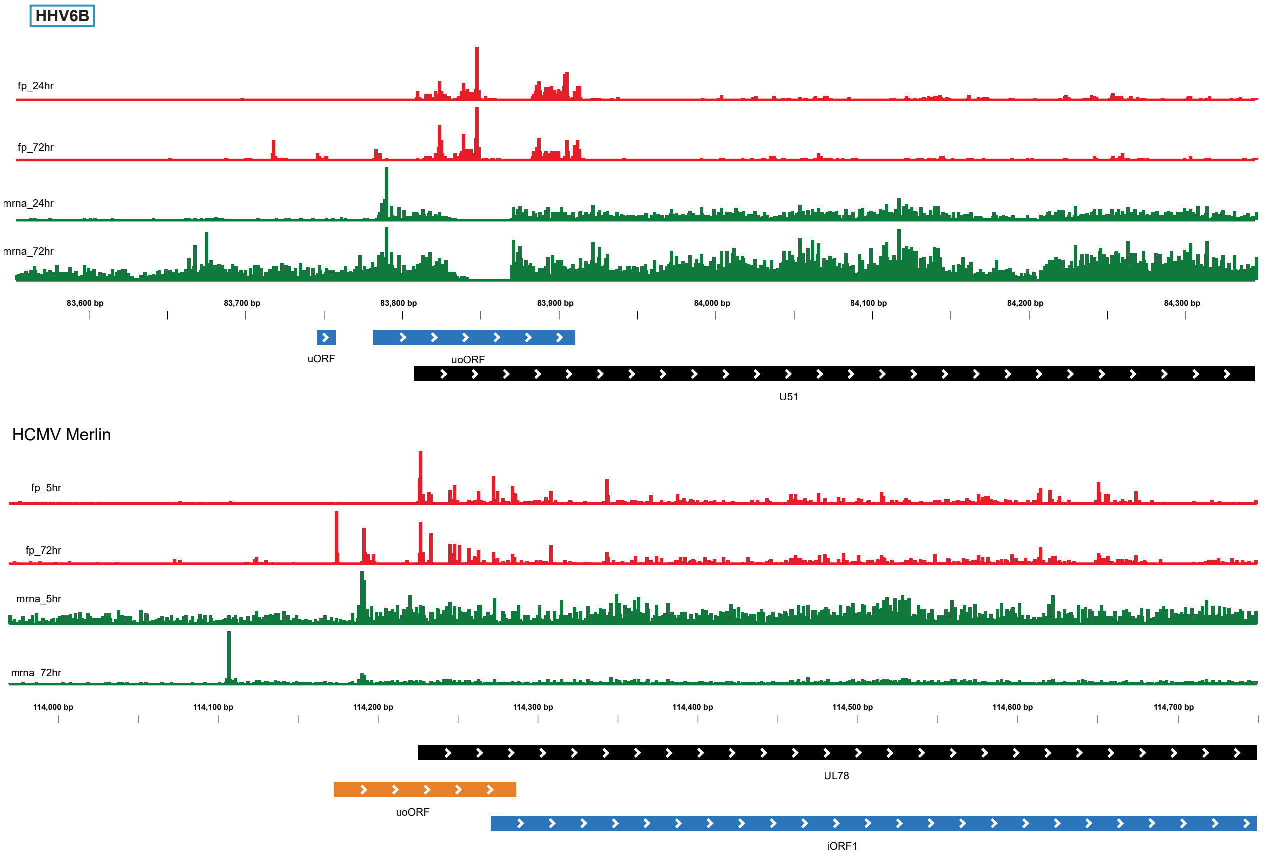

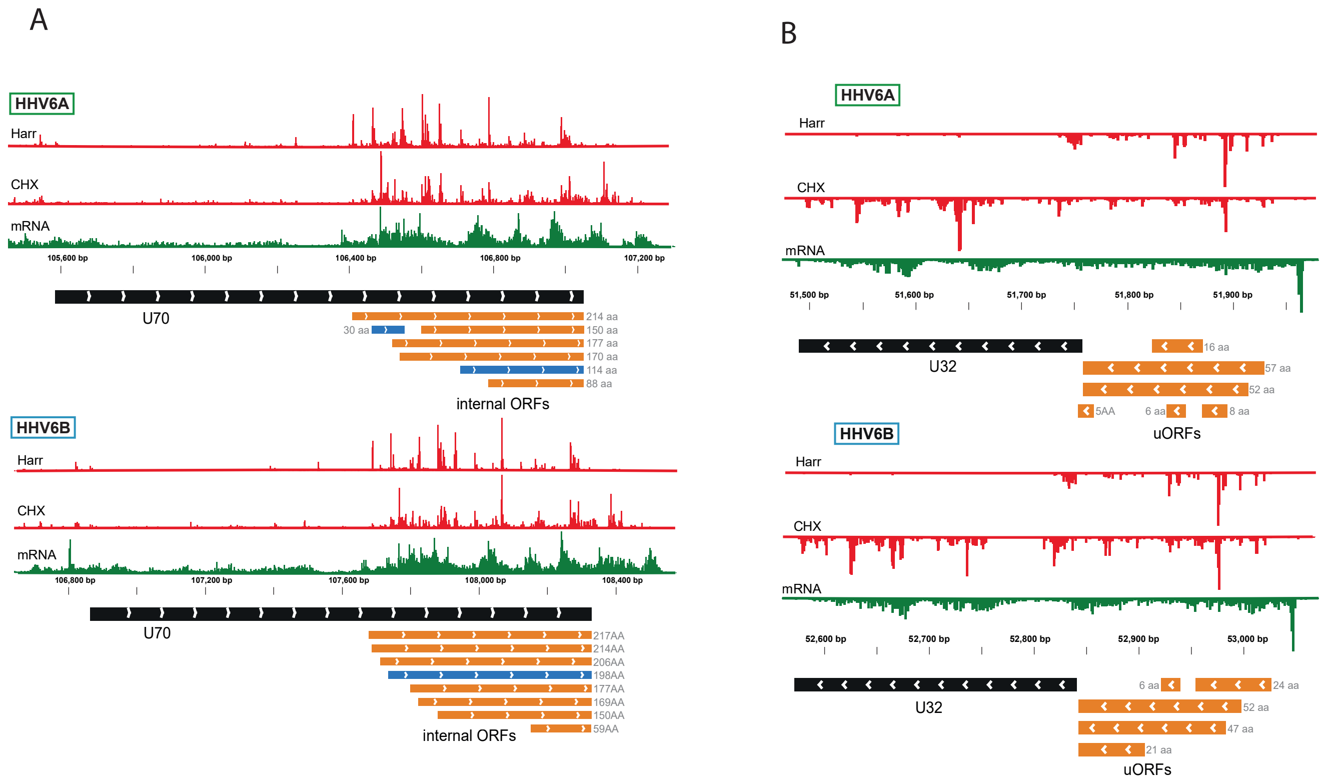

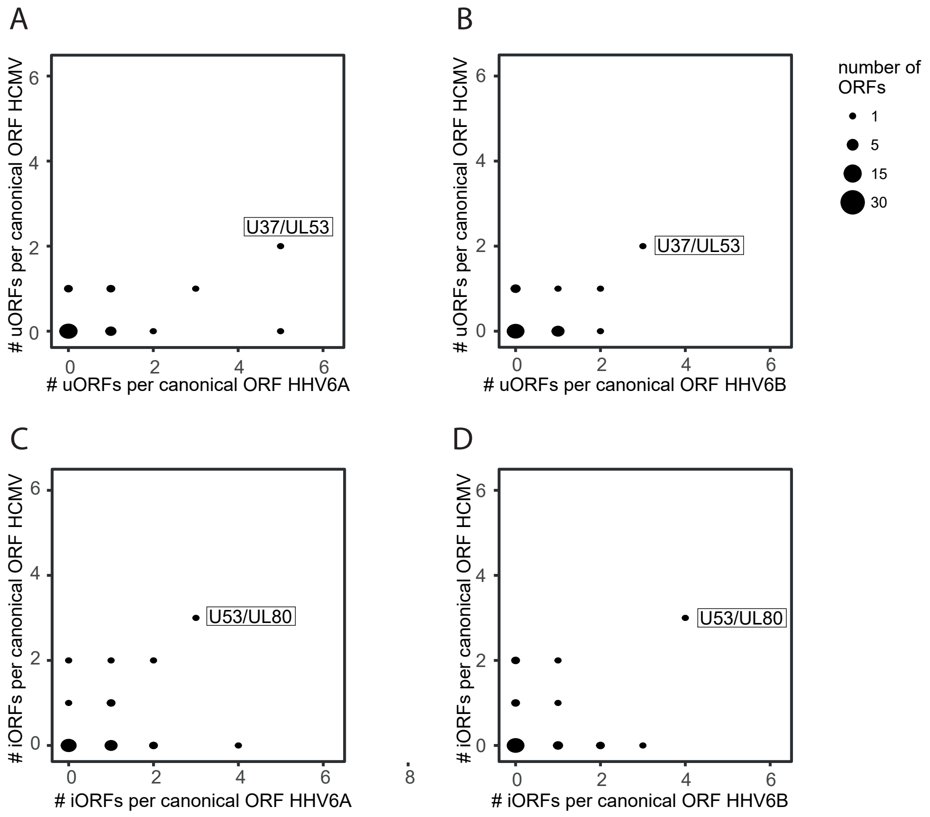
